# Single nuclear RNA sequencing shows altered microglial and astrocytic functions in post-mortem Parkinson’s disease tissue

**DOI:** 10.1101/2025.11.13.687853

**Authors:** Indra Roy, Rhalena A Thomas, Georgina Jimenez Ambriz, Sali Farhan, Valerio E.C. Piscopo, Thomas M Durcan

## Abstract

**Background:** Parkinson’s disease (PD) is a neurodegenerative disease marked by a progressive loss of dopaminergic neurons in the substantia nigra *pars compacta* (SNpc) and formation of misfolded protein aggregates. A growing body of research has implicated glial cell dysfunction in PD etiology, including the concentration of activated glial cells around protein aggregates in post-mortem tissue. Disruptions in the balance of pro– and anti-inflammatory immune response functions of the microglia and astrocytes is believed to contribute towards neurons being lost as the disease progresses. However, the molecular mechanisms remain unclear. To shed light on the role of inflammation in PD, this study analyses two public single nuclear RNA sequencing datasets of the SNpc from patient and control postmortem brain to identify altered molecular pathways in PD-associated microglia and astrocytes.

**Results:** The results show that both cell types have a significant upregulation in heat shock binding and misfolded protein response pathways, likely in response to the accumulation of protein aggregates. Microglia annotated with the MKI67 marker gene show a decreased expression in PD patient derived tissue. Markers associated with activated/reactive states in astrocytes and microglia are upregulated in PD samples. Notably, expression of genes associated with resting state microglia and non-inflammatory reactive state microglia are downregulated in PD microglia, including P2RY12, CSF1R, CSF2RA, CSF3R, and TGFBR1. Concurrently, genes associated with activated microglial states such as HSP90AB1 and GPNMB are upregulated. Among the top downregulated functions, genes associated with ion channel functions are downregulated in both astrocytes and microglia.

**Conclusions:** Taken together, the findings imply that astrocytes and microglia respond to protein misfolding pathology in PD by upregulating chaperone protein folding functions. Additionally, the profile of upregulated genes implies that pathways responding to oxidative stress are also activated. The downregulation of inflammation-associated genes in PD microglia paired with the upregulation of misfolding protein response pathways, suggests a switch from immune receptor functions to protein aggregate clearance by the end of disease stages. Finally, GPNMB emerged as a potential target for therapeutic intervention, as the one primary non-HSP gene that is significantly increased in PD-associated microglia.

**Abbreviations:** Parkinson’s disease (PD); neurodegenerative diseases (NDD); dopaminergic neurons (DNs); substantia nigra *pars compacta* (SNpc); microglia (Mg); Lewy body dementia (LBD); single nuclear RNA sequencing (snRNA-seq), single cell RNA sequencing (scRNA-seq); Gene Ontology (GO)

## 1.0 BACKGROUND

Several neurodegenerative diseases (NDDs) including Alzheimer’s disease (AD) and Parkinson’s disease (PD) are characterized as both proteinopathies – with aberrant post-transcriptional modifications and protein aggregates [1, 2], and neuroinflammatory disorders, with reactive glial cells [3–5]. Although the two mechanisms are hypothesized to overlap [5–8], the exact pathways by which they contribute to neurodegeneration remains unclear. A recent review suggested that glial cells maintain a balance of both pro– and anti-inflammatory functions in the brain and that disrupting this balance promotes the progressive loss of neurons observed across NDDs [9]. The inflammatory hypothesis suggests that dysregulated signaling pathways in glial cells contributes to neurodegeneration. Astrocytes, microglia, and invading T cells from the periphery together promote an inflammatory cascade that can be both pathogenic and protective in these NDDs [10]. We hypothesize that altered gene expression in patient-derived microglia and astrocytes will reveal altered molecular functions that are pathological drivers of proteinopathy, either in response to or concurrent with protein misfolding.

Studies exploring the role of glial involvement in NDDs have increased in recent decades. In AD, post-mortem tissue analyses consistently show increased reactive microglia (Iba-1^+^/HLA-DR^+^/CD11b^+^) around amyloid-β (Aβ) plaques [11]. A higher density of reactive microglia (HLA-DR^+^) is also observed in the substantia nigra *pars compacta* (*SNpc*) of PD patients in post-mortem tissue, particularly surrounding synuclein deposits [12]. Increased reactivity is observed in early disease stages through positron emission tomography (PET) imaging studies indicating potential causative or exacerbating roles of microglia and astrocytes. Radioligand-labelled TSPO, which marks primarily reactive microglia (and to some extent astrocytes) are increased in disease-relevant areas in both AD and PD; with symptom severity in both diseases correlating with microglial reactivity [13–15]. Recent studies in humanized AD animal models have suggested that microglial activation is responsible for spreading tau pathology through the brain, such that partially digested protein fragments are released through phagosomes [16]. Reactive astrocytes are also present in disease relevant areas from human post-mortem tissue in several NDDs, including the prefrontal cortex in AD, motor cortex in amyotrophic lateral sclerosis (ALS) and *SNpc* in PD [17]. Astrocytes cultured from LRRK2 mutant familial PD patient induced pluripotent stem cells (iPSCs) showed dysfunctional autophagy, increased reactivity, and impaired ability to phagocytose α-synuclein (α-syn) aggregates [18]. Similar activation of microglia is also triggered by neuronal α-syn through toll-like receptor 4 (TLR4), paired with upregulated autophagy receptors through nuclear factor kappa B (NF-κB) pathway in rat primary microglial cultures [19]. Thus, microglial reactivity under physiological conditions can be protective, clearing misfolded α-syn. On the other hand, α-syn induces pro-inflammatory cytokine secretion by iPSC-derived microglia [20]. A dose-dependent increase is seen in IL-1β and TNF in both iPSC-derived microglia [20] and mouse-derived primary microglia [21]. Thus, continued accumulation of misfolded protein aggregates may contribute to pathogenesis through chronic neuroinflammation. In microglia, membranous vesicles containing cytoplasm and released from the plasma membrane (exosomes) have been suggested to mediate prion-like transmission of synuclein to neurons [22]. A similar glia-to-neuron transmission of α-syn was observed in astrocyte-neuron coculture models [23]. Together this implies that glia influence the spreading of α-syn, thereby promoting neuronal cytotoxicity in PD and other NDDs [23, 24].

The mechanisms or pathways involved in PD pathology are still not well characterized. The literature suggests contradictory conclusions regarding the role of microglia in NDD [9, 25–27] Similar contradictory conclusions are also found regarding the role of reactive astrocytes in NDD. Partly, this is due to the heterogeneity of phenotypes expressed by both glial types. Earlier researchers attempted to categorize astrocytes and microglia as “reactive” or inflammatory while categorizing alternately reactive states as “anti-inflammatory”. This conceptualization has become increasingly contentious as the complexity of individual cytokines’ activity are revealed to be more context-dependent; for example, IL-10 is often referred to as an anti-inflammatory cytokine since it inhibits pro-inflammatory cytokines TNF-alpha, IL-1β, and IL-6. However, prolonged activation of IL-10 can also trigger a resurgent increase in IL-1β and IL-6, thus indirectly contributing to pro-inflammatory processes. Furthermore, newer technologies such as single cell RNA sequencing have revealed that both astrocytes and microglia can display a variety of activation states that are challenging to categorize within a spectrum of reactivity. This makes it more difficult to identify the functional contribution of glial cells to PD pathogenesis.

Here, we attempt to delineate the functions altered across heterogenous populations of microglia and astrocytes by analysing two publicly available single nuclear RNA sequencing (snRNAseq) datasets from human post-mortem midbrain tissue of idiopathic PD patients and age-matched controls [28, 29]. Kamath et al. enriched their dataset for dopaminergic neurons to identify vulnerable subtypes of DA neurons in PD and Lewy body dementia. Extracting microglia and astrocytes from their original dataset, we focus specifically on PD samples and perform differential gene expression (DGE) analysis and subsequently pathway analysis to identify whether genes associated with certain molecular functional pathways are altered in PD microglia and astrocytes compared to control samples (28). While the larger dataset (28) is used as the primary focus of our analysis, a secondary dataset is also analysed to identify conserved signatures of altered microglial and astrocytic functions (29).

### 2.0 METHODS

### 1. Data acquisition and processing

For the Kamath [28] dataset, we obtained the filtered gene expression matrix for the complete dataset from the Broad Institute’s Single Cell Portal, and extracted cells annotated as astrocyte or microglia in separate data objects. For quality control purposes, Kamath et al removed cells with >10% of reads of mitochondrial genes and a significantly lower or significantly higher UMI compared to the rest of the dataset. The cell type and cell subtype annotations provided by Kamath in a metadata file were used in this study. Although they grouped macrophages as a subtype under the broad categorization of microglia, we excluded them and analysed the remaining cells under microglia. Similarly, under astrocytes, we removed cells annotated as ependymal cells and analysed the remaining cells as astrocytes. The original dataset includes samples from patients with Lewy Body Dementia, which were excluded from our analysis. **Figure 1** shows UMAPs of the original dataset, highlighting the cells that were subsequently extracted as individual microglia and astrocyte data objects. UMAPs of the resulting data objects with included subtype labels are shown in **Figures 2A and 2C**.

**Figure 1.**
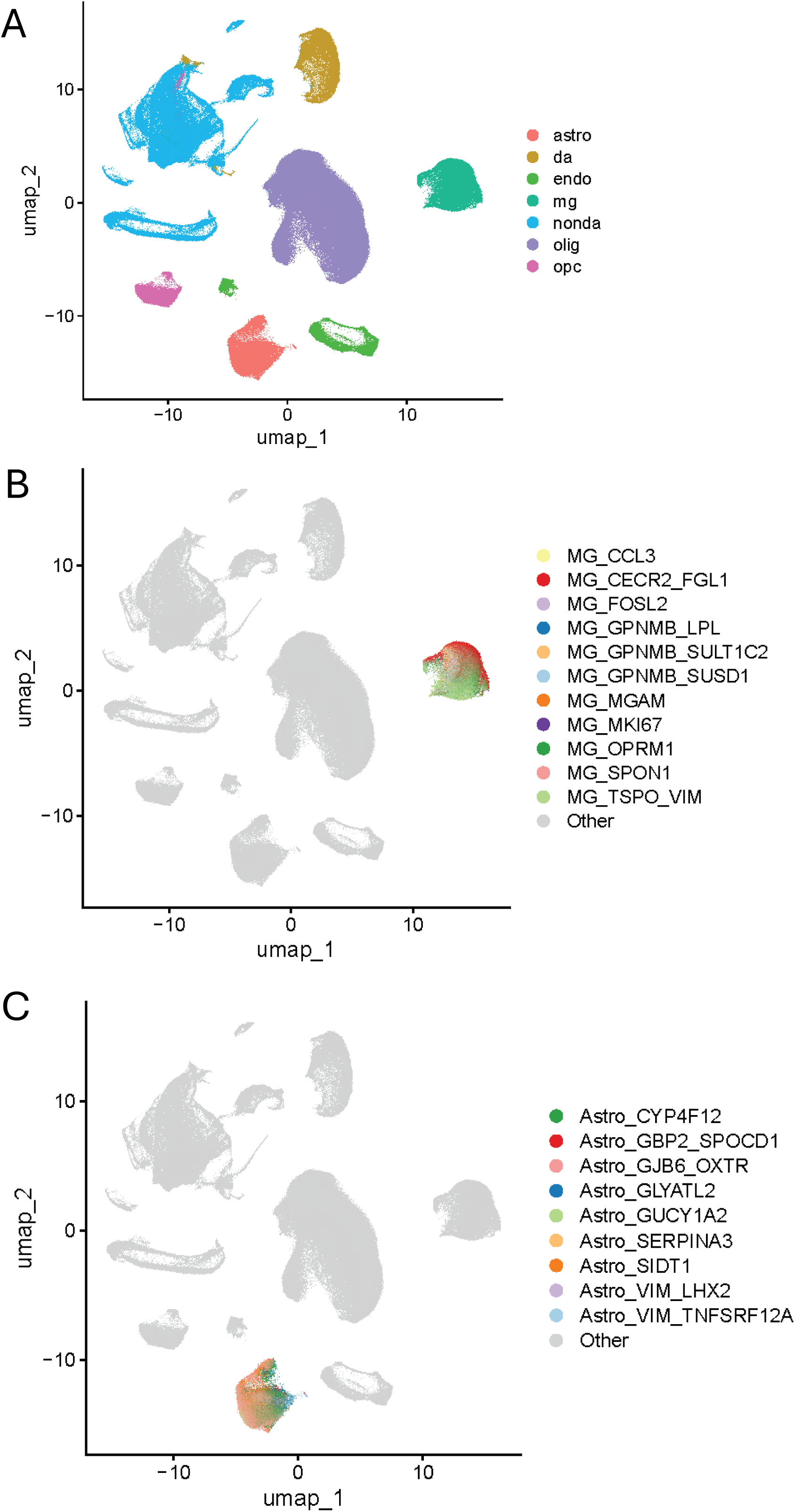
Cell type distribution of all human midbrain cells in the Kamath dataset. (A) Complete UMAP of cell type distribution with contracted annotation that were used for analysis and subsetting. (B) Microglial cluster highlighted and labelled with their original subtype names given by Kamath et al. (C) Astrocyte cluster highlighted and labelled with their original subtype names given by Kamath et al.

**Figure 2.**
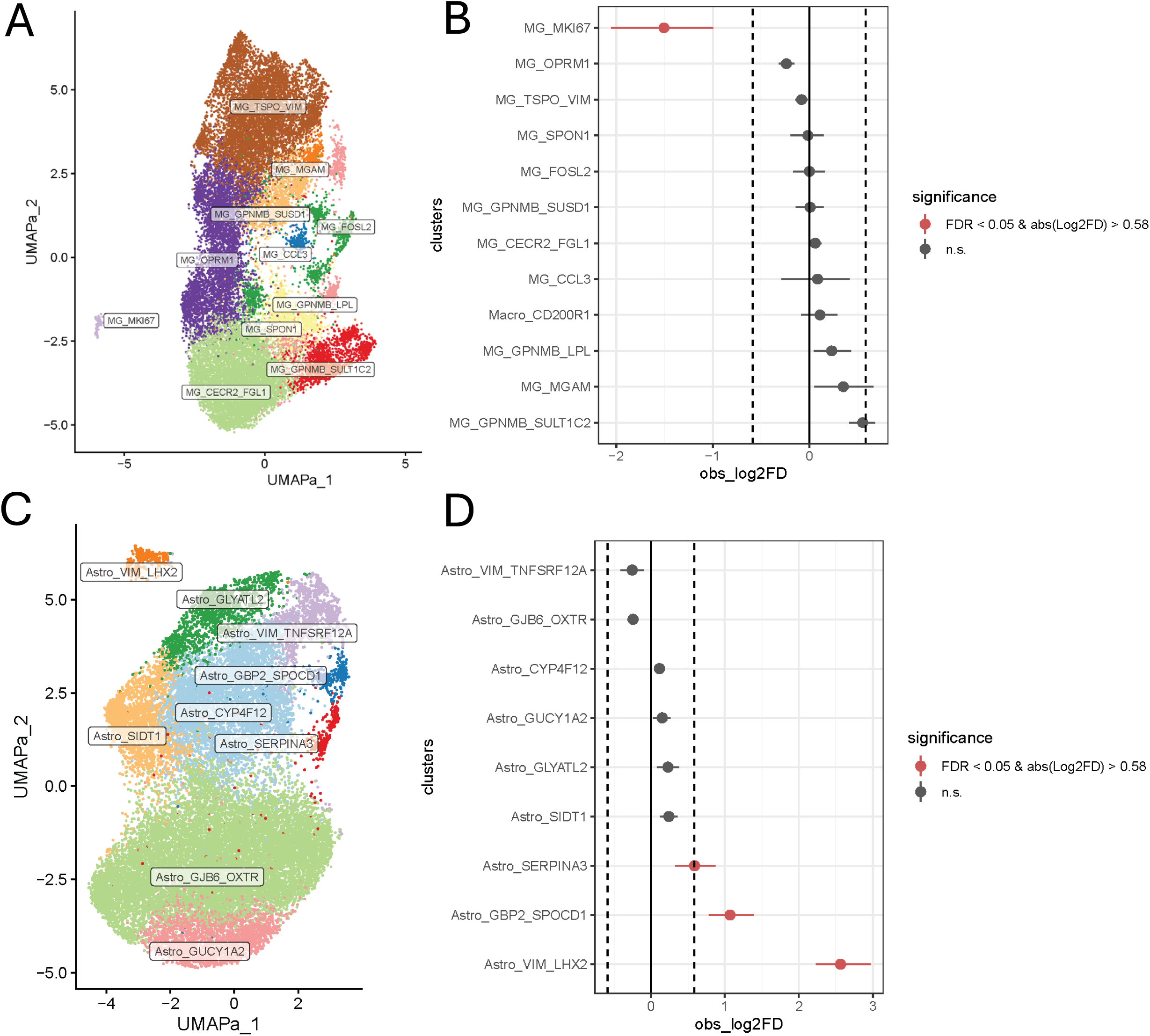
Distribution of selected cell subtypes in extracted microglia and astrocyte objects. (A) Subtypes annotated as microglia retained for subsequent analyses includes only those derived from PD and control donors and excludes macrophage cells. (B) Proportional representations of each annotated microglial subtype in PD as compared to their expected proportions based on control donors. (C) Subtypes annotated as astrocytes retained for subsequent analyses includes only those derived from PD and control donors and excludes ependymal cells. (D) Proportional representations of each annotated astrocyte subtype in PD as compared to their expected proportions based on control donors.

For the Smajić et al. [29] dataset, the raw data was downloaded from Gene Expression Omnibus (GEO; GSE157783). The raw sequencing data was processing using the scRNAbox pipeline [30]. FASTQ files were aligned to the human genome (GRCh38), and the expression count matrix, feature and single cell barcode files were generated using CellRanger v.7. Putative cells were filtered to retain cells with a minimum of 1000 unique gene transcripts per cell and less than 25% mitochondrial RNA. Individual samples were integrated using the Seurat V4 integration functions [30]. Cells from the integrated data object were subsequently clustered: the top 3000 most variable features were input into a principal component analysis dimensional reduction. The top 30 principal components were used as the input for generating a shared nearest neighbour graph (SNN) and uniform manifold approximation and projection (UMAP) visualization. Clusters were identified using the Leiden network detection algorithm with the SNN graph as the input and run for a range of clustering resolutions, with 0.2 for annotation selected as the resolution. Broad cell type annotations were applied using step7a of the toolbox, with microglia and astrocytes subsetted into separate objects.

### 2. Cell population testing

A proportionality test was performed to identify any contributions of disease ontology to cell type numbers. The function ScProportion [31] was used to perform permutations tests comparing the proportions of each cell type within the PD group against the control group compared to the distribution of proportions generated by randomly shuffled labels between PD and control for each cell type. ScProportion, a permutation test was used to compare the proportions of cell types within the PD group against the proportions of cell types in the control group.

### 3. Differential gene expression (DGE) analysis

Differential gene expression (DGE) analysis was performed using Seurat (v4.3.1) *FindMarkers* function contrasting PD against control, using the MAST method [32], with a p-value threshold of 0.05. DGE analysis was performed separately for all cells in the astrocytes and microglia datasets. This analysis identifies genes that are expressed at significantly higher or lower levels in the experimental group compared to control. The function outputs a gene list accompanied by the log-fold change of each gene relative to their expression in control cells.

### 4. Molecular function analysis

Gene set enrichment analysis (GSEA) was performed on the log fold change (logFC) values of the DGE gene lists for enrichment of the Gene Ontology (GO) molecular function terms. Using the R package ClusterProfiler’s gseGO function, the logFC is compared across differentially expressed genes to rank up and down-regulated molecular functions. This analysis identifies whether genes associated with specific molecular functions are enriched in PD in an upward or downward direction.

## 3.0 RESULTS

### Microglia and astrocyte populations differ in vulnerability and proliferation

One way to determine whether cell types are impacted in disease is to perform a proportionality test that identifies whether certain cell types are present at higher or lower levels in the experimental group than their proportion in the control group. A lower than predicted cell number can potentially indicate if a cell type is vulnerable and thus selectively lost in a disease. A higher than predicted number would suggest that the cells in question are upregulated in disease states either pathologically or in response. In the Kamath dataset [28], a comparison of total cell populations between PD and control showed that cell numbers were within the expected range for all cells except dopaminergic neurons which were decreased in PD compared to their proportion in control patients (**Supplementary Fig. S1**), in line with previously published findings for this dataset [28].

Within the eleven microglia subsets, a small subgroup characterized by the cell proliferation marker MKI67 expression (Mg_MKI67) were reduced in the PD cohort, revealing a subtle microglial vulnerability (log^2^FDR ≈ –1.8) in this microglia subtype (**Fig. 2B**). The protein Ki67 encoded by MKI67 is involved in the regulation of mitotic nuclear division. This potentially indicates that the proportion of proliferative microglia are decreased in the final disease stages of PD, although the overall population of microglia are unaffected. In contrast, there are three subtypes of astrocytes that show increased proportion in PD patients: Astro_VIM_LHX2 (log^2^FDR ≈ 2.6), Astro_GBP2_SPOCD1 (log^2^FDR ≈ 1.2), and Astro_SERPINA3 (log^2^FDR ≈ 0.6) (**Fig. 2D**). The marker genes they are annotated by are primarily associated with inflammatory processes (LHX2, SERPINA3 and GBP2), or with broad cellular functions including cell migration and attachment, and structural stability (VIM and SPOCD1).

### Chaperone protein binding is strongly upregulated in PD microglia and astrocytes

Given that the proportion of microglia and astrocytes as broad cell types are not altered between PD and control patients, we sought to analyze differential gene expression (DGE) between PD and control in microglia and astrocytes as separate data objects. The resulting gene lists were used for gene set enrichment analysis to characterize molecular functions that were enriched in PD cells compared to control.

PD is primarily a synucleinopathy characterized by protein misfolding, although recent studies have pointed to inflammation and lipid dysfunction as pathways also implicated in the development of PD [33]. Our analysis shows that both astrocytes and microglia play a role in responding to pathological protein misfolding. Heat shock proteins such as HSP90AB1, HSP90AA1, DNAJA4 (an HSP40 variant), and co-chaperone protein BAG3 are all among the top 20 upregulated genes in both astrocytes and microglia, with the largest log fold change (**Fig. 3 and 4**). In both cell types, heat shock protein binding (GO: 0031072) and protein folding chaperone (GO: 0044183) are among the top 10 upregulated GO molecular function terms in PD patients (**Fig. 5A and 6A**). GeneRatios between 0.7-0.8 for these functions indicates that 70-80% of known genes associated with this function are enriched in PD microglia. ATPase regulator activity is also among the most significantly upregulated functions, implying that energetic needs may drive microglia-mediated protein folding functions as a disease response (**Fig. 5A**). The same trend is also observed in PD astrocytes with enriched gene expression associated with ATPase regulator activity (**Fig. 6A**), albeit with a smaller gene count ratio.

**Figure 3.**
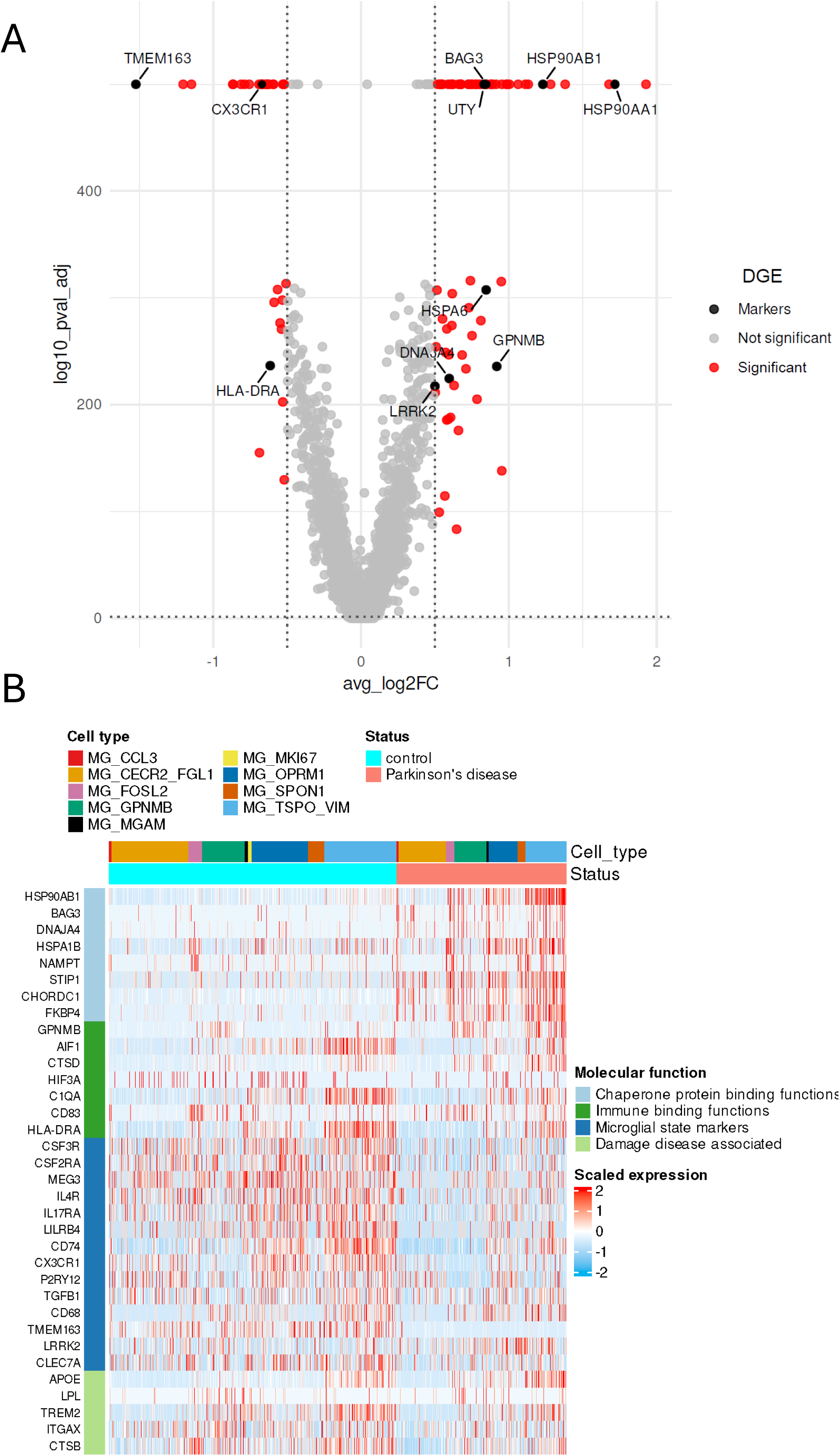
Differential gene expression in PD microglia highlighting selected disease-salient genes. (A) Volcano plot demonstrates altered gene expression in PD microglia; genes that were among the top 20 altered genes by p-value that also had a log fold change ±0.5 are labelled. (B) Heatmap expression of disease-relevant genes in PD microglia shows an increase in several prominent genes associated with chaperone protein binding functions, and a downregulation of key genes associated with immune binding functions, microglial state markers, as well as some damage associated markers.

**Figure 4.**
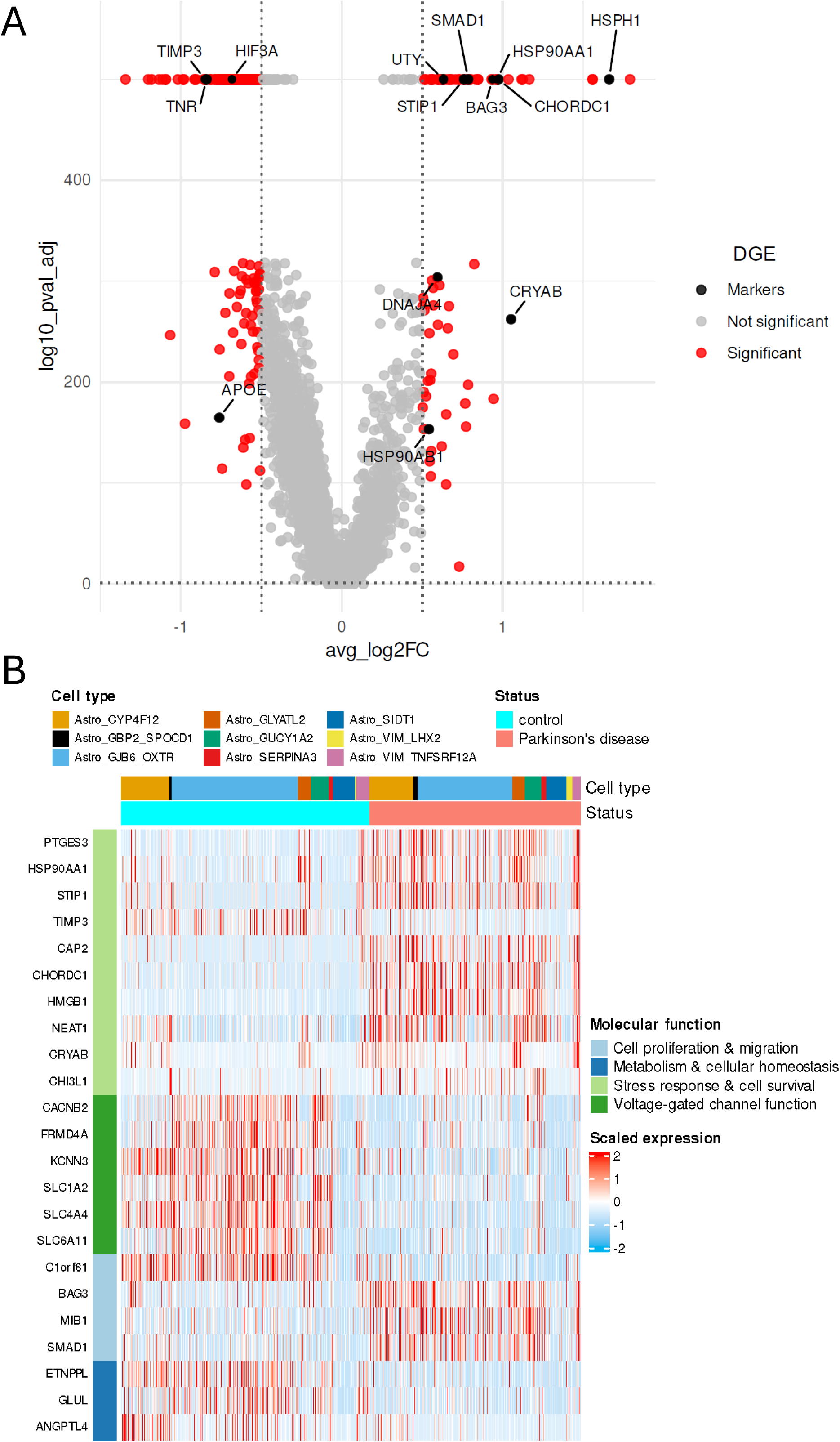
Differential gene expression in PD astrocytes highlighting selected disease-salient genes. (A) Volcano plot demonstrates altered gene expression in PD astrocytes; genes that were among the top 20 altered genes by p-value that also had a log fold change ±0.5 are labelled. (B) Heatmap expression of disease-relevant genes in PD astrocytes shows an increase in genes associated with stress response and cell survival, and cell proliferation and migration, as well as a decrease in genes associated with metabolism and cellular homeostasis, and voltage-gated channel functions. Where a gene is associated with more than one function (BAG3), it is mapped in the category with fewer genes.

**Figure 5.**
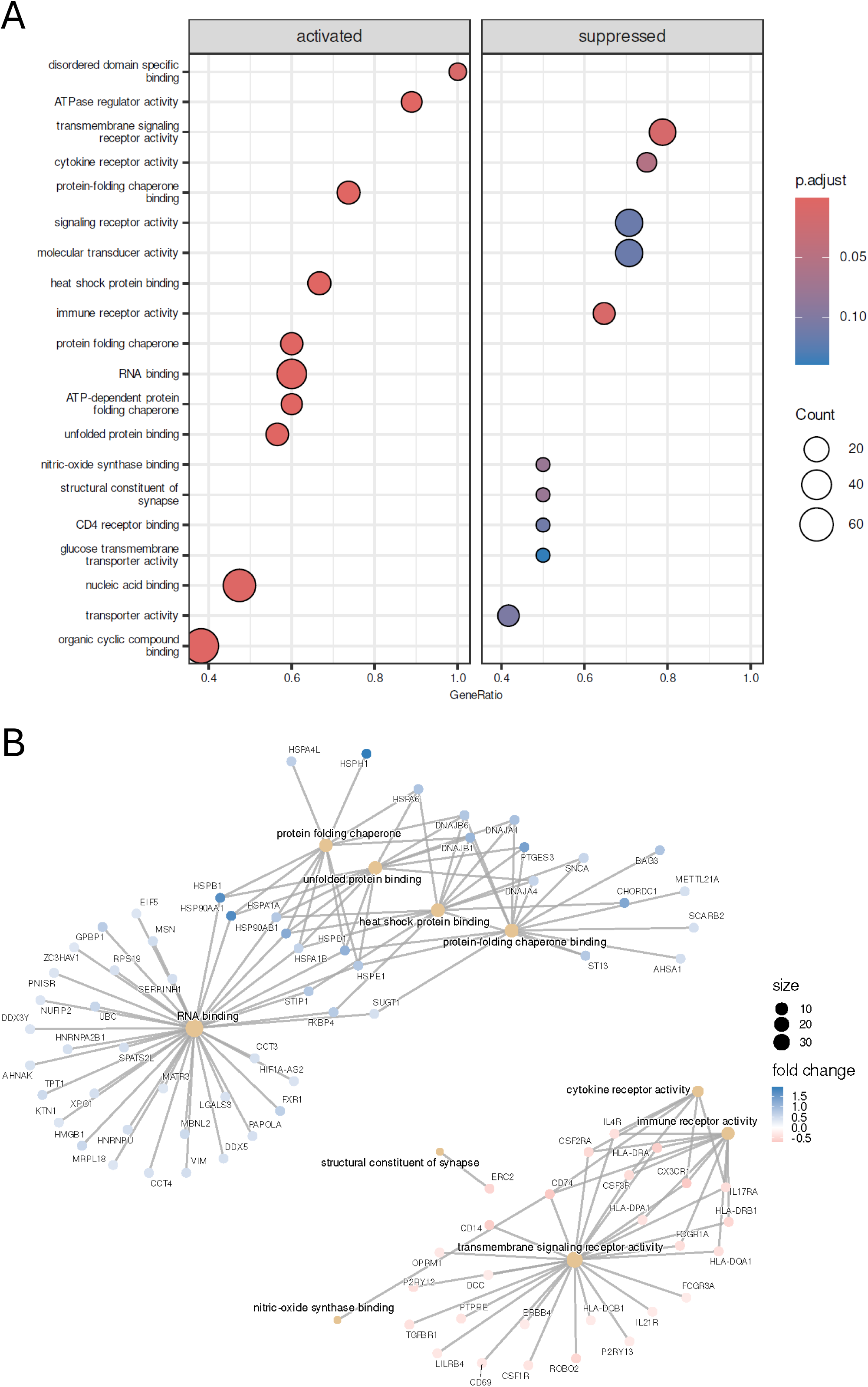
Molecular function enrichment in PD microglia. (A) Dot plot graph of the top 10 most upregulated and downregulated functions shows the gene count that are in that enriched function, as well as the GeneRatio that shows the proportion of known genes associated with that function that are also enriched on this analysis. Several GO terms associated with chaperone protein binding functions are in the activated column, along with ATPase regulator activity. On the other hand, transmembrane signalling activity, specifically cytokine and immune receptor activity are in the suppressed column. (B) The top 5 most upregulated and downregulated functions and the genes associated with this enrichment is shown as a CNet plot that also shows relationships between enriched genes that are represented in more than one function.

**Figure 6.**
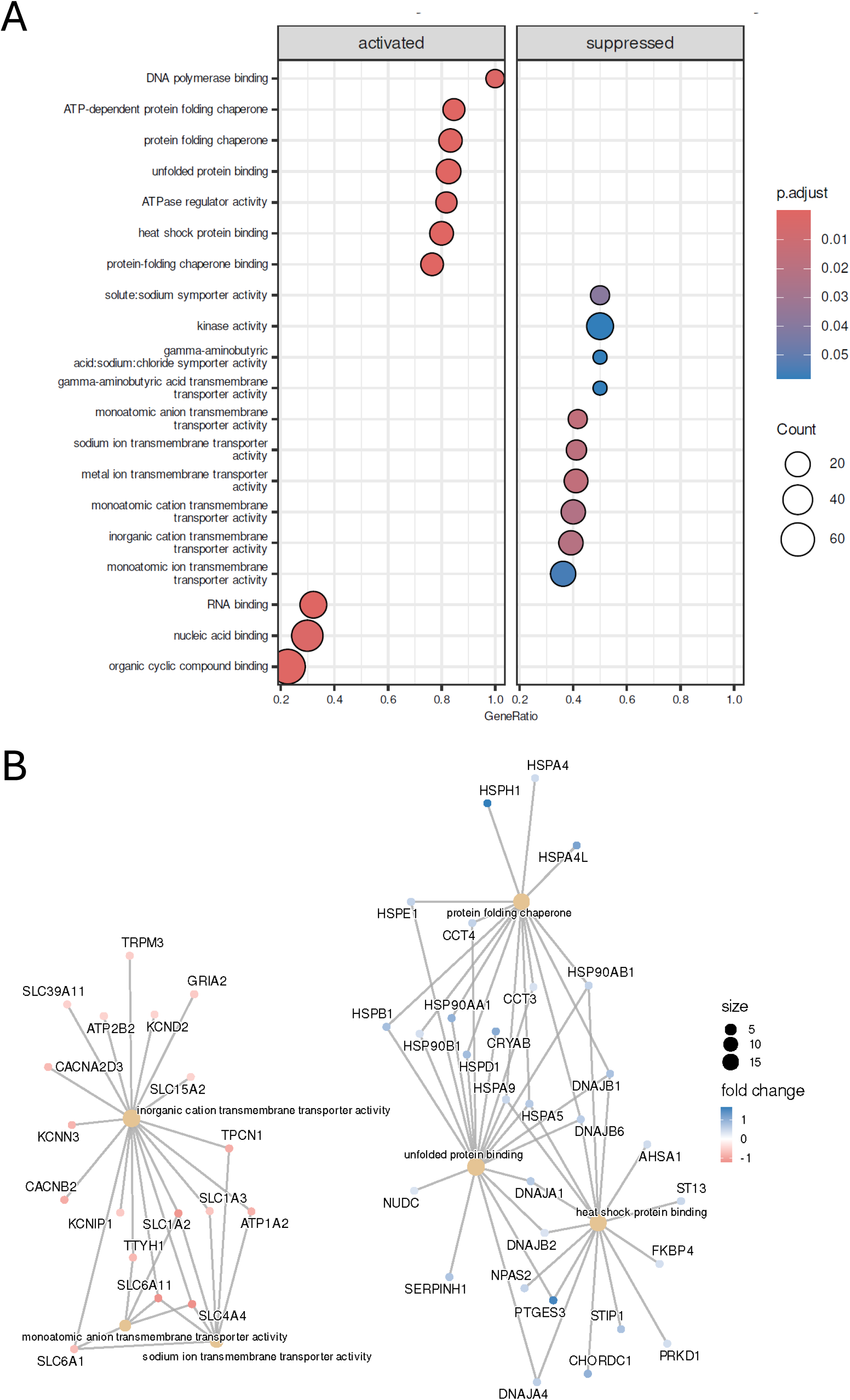
Molecular function enrichment in PD astrocytes. (A) Dot plot graph of the top 10 most upregulated and downregulated functions shows the gene count that are in that enriched function, as well as the GeneRatio that shows the proportion of known genes associated with that function that are also enriched on this analysis. Several GO terms associated with chaperone protein binding functions are in the activated column, In the suppressed column, most of the GO terms are associated with voltage gated or ionotropic receptor transmembrane transporter activity.(B) The top 5 most upregulated and downregulated functions and the genes associated with this enrichment is shown as a CNet plot that also shows relationships between enriched genes that are represented in more than one function.

In both astrocytes and microglia, the activation of chaperone protein binding is enriched by an increase in chaperone proteins (such as STIP1/STI1) and heat shock proteins (HSPs) themselves (**Fig. 5B and 6B**). A range of HSPs are upregulated in both, with the strongest upregulation in known inflammation markers HSP90AB1 & HSP90AA1 (**Fig. 3A and 4A; Supplementary Tables 1 and 2**). Hsp70 binding is also implicated, as is a range of DNAJC17 (also known as HSP40) variants. As well, in mouse microglial cells, BAG3 was shown to promote clearance of α-synuclein fibrils and autophagy in an inflammatory environment while suppressing NLRP3 inflammasome formation [34, 35]. This suggests that a response to misfolded protein accumulation in PD may be a key process by which microglia mediate later disease stages. The increased expression of these genes supports the hypothesis that microglia, and to some extent astrocytes, respond to the accumulation of protein aggregates by upregulating protein folding pathways.

### Immune receptor functions are impaired in PD microglia, accompanied by an increase in oxidoreductase functions

Often an increase in HSPs and the associated chaperone folding genes’ expression is seen as evidence of increased cellular stress. Previous studies posited that increased glial activation occurs early in PD, with several PET studies showing increased TSPO-radio-ligand labeled cellular activity in recently diagnosed PD patients [13, 14]. Cell culture studies have also shown an increased secretion of pro-inflammatory cytokine in response to synuclein fibrils, through NLRP3 and TLR4 mediated activation of cultured microglia from various sources, including rat and mouse primary microglia and human-iPSC-derived microglia [19–22]. In this analysis we had the opportunity to examine altered immune function associated pathways at the end of the disease, and the results imply that pro-inflammatory reactivation by microglia may be a physiological response lost over the course of prolonged proteinopathy, significantly downregulated compared to age matched control tissue donors.

Several of the top 10 most downregulated functions in PD microglia are associated with receptor activities, including immune receptor activity (GO: 0140375) (NES=-2.36, adj.p.val=0.01) (**Fig. 5A**). The cnet plot shows that immune receptor activity is among the top 5 downregulated functions and is enriched by HLA genes and several interleukin receptor genes (IL-4R, IL-17R, IL-21R) (**Fig. 5B**). While IL-4 is generally considered to be anti-inflammatory, IL-17 is considered pro-inflammatory, produced primarily by helper T-cells and associated with chronic inflammatory conditions. IL-21 can be considered both pro– and anti-inflammatory depending on the context; its expression is triggered by CD4+ cells and can enhance inflammatory responses, but it can also promote the proliferation of regulatory T-cells that attenuate pro-inflammatory responses.

Heatmaps also highlight a decrease in the expression of several microglial markers that are associated with motile branching microglia in PD (**Fig. 3B**). HLA-DRA, CX3CR1, and colony stimulating factor genes CSF2RA and CSF3R are strongly expressed in control microglia, and a decreased expression of these genes in PD microglia are suggestive of a loss or decrease in surveying, quiescent microglia. Concurrently, the commonly known PD-risk-associated gene LRRK2 is also upregulated in PD microglia (**Fig. 3A**); with mutations in LRRK2 known to contribute towards PD pathogenesis through a myriad of ways by disrupting vesicular formation, cellular trafficking, lysosomal and autophagic dysfunction, combined with increased pro-inflammatory cytokine secretion [36–39]. The downregulation of genes associated with homeostatic microglia as well as genes associated with pro-inflammatory pathways imply a broad downregulation of all microglial immune functions in end-stage PD. Microglia may also become exhausted in later stages and be unable to keep up with their own regeneration, which would align with the decreased expression of microglia marked by the proliferation marker MKI67 (**Fig. 2B**).

### Transmembrane receptor functions associated with quiescent and non-inflammatory activated microglia are downregulated in PD

Our observations on immune receptor activity described above aligns with a lower branch of the GO term tree in the form of transmembrane signalling receptor activity (GO: 0004888), which is another top downregulated function in PD microglia (**Fig. 5A**). In addition to the immune receptor genes, this enrichment includes genes that promote microglial surveillance, motility, and response to pattern or damage associated markers (PAM/DAMs), such as P2RY12, CSF1R, CSF2RA, CSF3R and TGFBR1 (**Fig. 3B**). The downregulation of these genes, particularly receptors for colony stimulating factors indicates a reduced ability of microglia to respond to damage and inflammation in PD. Recent studies indicate that P2RY12 is differentially expressed at higher levels in resting and non-inflammatory activated microglia [40, 41], with further evidence implying that microglial chemotaxis occurs in response to neuronal injury. Thus, our analysis supports the notion that microglial homeostasis is disrupted in PD, particularly affecting the different microglial states responsible for damage response and resolution functions. One notable exception to this pattern is the increased expression of GPNMB in PD microglia across both datasets, which is typically associated with anti-inflammatory response in macrophage-like cells. This could indicate that earlier attempts at damage-resolution from responding microglial cells may simply become ineffective or inadequate over time, despite increased transcription at the RNA level.

### Voltage gated ion channel dysfunction in PD astrocytes and microglia

Astrocytes are thought to play a role in regulating neural cell signalling and synaptic transmission, through their extensive network of neurotransmitter channels (such as glutamate and GABA) and voltage gated ion channels (such as Ca2+, Na+, and K+ channels) [42]. In PD animal and cell culture models, altered characteristics of several astrocytic ion channels have also been associated with pathological or neurodegenerative phenotypes [43]. When it comes to voltage gated ion channel activities in PD astrocytes, the two datasets contradict each other. Functional terms associated with these are downregulated in the Kamath astrocytes and includes Na+/K+/Ca2+ channel constituents, regulators and binding functions indicating a broad impairment of signalling and channel functions in PD astrocytes (**Supplementary Table 3**). While a similar effect is seen in the Smajić dataset regarding a decrease in Ca2+: Na+ antiporter activity, only one gene, SLC24A2, is associated with this function (**Supplementary Table 4)**.

Alternatively, ion channel dysregulation could be an effect of mitochondrial dysfunction, as many voltage-gated channel functions are ATP-dependent, and ATPase-related functions are also impaired in astrocytes and microglia. Energy requirements may be overtaken by protein folding and chaperone protein binding functions that are upregulated in both microglia and astrocytes in PD, thus leaving normal channel functions impaired. Summarily, in end stage PD patients, increased astrocytic response towards managing proteinopathy may lead to a compromised ability to perform normal channel functions either mechanistically due to changes in membrane permeability or gradient potential, or energetically due to ATPase-related mitochondrial dysfunction.

### Validation of findings with chaperone protein binding in a second dataset

In a second snRNA sequencing dataset, the strongest replicated finding was the enrichment of protein folding chaperone and heat shock protein binding functions with an increased expression of key HSPs such as HSP90AA1, HSP90AB1, DNAJB1, and STIP1 (**Supplementary fig. 2**). This effect remained consistent in both microglia and astrocytes (**Supplementary fig. 3**), despite the smaller total number of genes with altered DGE for both cell types in this dataset. In this dataset we did not see altered DGE specifically for cytokine receptor binding or immune receptor activity as broad GO molecular functional terms. However, isolated genes associated with immune functions, such as IL1B, TGFB1, and CD83 are all increased albeit with a very small log fold change (**Supplementary table 5**). As in the Kamath dataset, LRRK2 expression is also increased in PD microglia. Most notably, GPNMB remains one of the few non-HSP genes to be significantly upregulated in PD microglia (**Supplementary Fig. 2A**); GWAS studies have shown GPNMB is among one of the top risk genes associated with PD [44–46], making it an excellent candidate gene for studying the impact of its altered expression on microglial function in experimental models of PD.

## 4.0 DISCUSSION

Inflammation is a known pathological feature of numerous neurodegenerative disorders including PD, and its interaction with proteinopathy (e.g., protein misfolding or aggregation) has been characterized with limited success and mixed results in cell cultures and animal models. In this study, we leveraged two large public datasets containing single cell RNA sequencing data from human post-mortem brain tissue to investigate altered molecular functional signatures in two main glial cell types in PD. While the original studies focused on dopaminergic neurons that are the most affected cells in PD, we sought to understand whether there were underlying alterations in microglia and astrocytes to explore additional pathological degenerative mechanisms.

Our findings show that heat shock protein binding and misfolded protein binding functions were among the most upregulated functions. This is characterized by an increased expression of HSPs and chaperone binding proteins such as STIP1/STI1. Overexpression of STIP1 has been shown to aggravate α-synuclein-mediated toxicity in a recent mouse model [47]. This supports earlier hypotheses that microglia attempt to clear misfolded protein aggregates in response to proteinopathies including in PD [19–21, 48]. Notably in the Kamath dataset, the increase in heat shock protein binding functions is accompanied by a decrease in immune receptor functions in microglia shown by a significant downregulation of HLA and interleukin receptor genes. While key markers of glial activation (HSP90AB1, GPNMB, IL-1B) continue to be present and upregulated in PD microglia, it is accompanied by a downregulation of genes associated with resting or surveillant microglia (P2RY12, CSF receptor genes, and TGFBR1). Although GPNMB is normally associated with pathways that ameliorate pro-inflammatory signaling [49–51], one study demonstrated a potential role for it in triggering intracellular processes required for pro-activation signalling [52], thus suggestive of a dual role for GPNMB in mediating complex microglia states. The importance of this gene as a potential focus for future investigation is further emphasized by the fact that it is one of the few non-HSP gene that is upregulated in PD microglia across both datasets.

For several years, debate surrounding the spectrum of glial activation has argued against a two-dimensional view that posits glia as “at rest” on one end and “activated” at the other [53]. Our findings provide further support for the idea that “activated” microglial states need not necessarily include pro-inflammatory features. Rather, activated microglia may have a range of activation states where different functions are prioritized in response to their environment. Microglia marked by proliferation gene MKI67 show a decreased expression in PD, hinting that physiological levels of proliferation are not maintained in PD. Given that post-mortem tissue constitutes an end-stage of PD; combined with the finding that misfolded protein binding functions are upregulated, we propose that the energy requirement for the latter forces pathological microglia switch from their physiological PAM/DAM response functions.

The prominence of genes associated with protein misfolding functions in both microglia and astrocytes of PD patients support the idea of glial involvement in attempting to clear aggregates. This process is likely to be energy intensive, which is validated by the increased expression of genes associated with ATPase regulation and ATP dependant activity. This is also supported by the decreased expression of the specific microglia subtype characterized by a proliferation marker MK167 (Mg_MK167). Mitochondrial dysfunction resulting from excessive energy expenditure could account for the loss of homeostatic immune receptor function of microglia [54]. Thus, while there is no significant change in the total number of microglia and astrocytes, their efficacy is compromised. This, combined hand in hand with glial transmission of fragmented proteins across the brain, increased activation in pre-clinical stages, and channel dysfunction that compromises the integrity of blood-brain barrier paint astrocytes and microglia in roles of prominent pathogenic drivers of PD.

The findings regarding genes associated with inflammation may appear conflicting or inconclusive between the two datasets. As mentioned before, while the size of the Kamath dataset allowed some discrimination between PD and control microglia in characterizing differential gene expression of notable inflammation associated genes such as CD74, HLA-DRA, AIF1 and C1QA, a similar effect was not observable in the Smajić dataset due to a broad upregulation of these genes across both donor groups. A recent publication from the Macosko group showed that microglial cells in particular were vulnerable to ex-vivo transcriptional alterations, dependant on several key experimental variables such as tissue dissociation protocol, cell isolation technique, brain regions, and the use of different scRNA-seq techniques [55]. Additionally, the very nature of single *nuclear* sequencing datasets excludes markers that would be present in cytosolic components, which account for most acute inflammatory functions. In future research, as single cell sequencing technology becomes more accurate and accessible for preserved tissues combined with advances in spatial transcriptomics, a more complete picture of transcriptomic changes in pathological glial cells will be possible. In the meantime, these results hint towards affected immune pathways such as those involving C1QA and TGFBR1 that can be investigated in *in vitro* models in earlier disease stages.

Post-mortem patient data analysis examines tissue at the end stages of disease and therefore, the end stages of glial dysfunction. Microglia and astrocytes are dynamic cell types with a range of activation states, and an end disease model may only capture the fatigued state of the cells following prolonged activation. Cell culture studies have shown that there is an efficiency curve for microglial clearance of protein aggregates, where at initial stages, it is a protective mechanism [48]. However, with increased accumulation and a decline in phagocytotic efficacy, microglia begin releasing fragmented misfolded proteins leading to a propagation of proteinopathy in a prion-like fashion [15, 16, 22]. A similar role in spreading misfolded protein fragments has also been posited in astrocytes [56]. In astrocytes, altered gene expression of voltage gated channel functions indicate their role in mediating altered synaptic transmission, likely impacting their communication with other cell types including microglia and neurons. Reactive states of microglia and astrocytes have been shown to impact each other [17], possibly creating a feedback loop of reactivation that exacerbates neurodegeneration.

Treating these observations as a limited end-stage disease model allows us to reconcile these finding with earlier in-patient findings. Positron emission tomography (PET) in PD patients have shown increase activation of radio-ligand labeled TSPO+ microglia in a sample of PD patients with a range of disease progression measured by the UPDRS; they found no correlation between disease stage and microglial activation [14], suggesting that altered states likely precede external diagnostic symptoms of PD. Similar findings in AD [13] and existing review literature [26, 57] suggests that the inflammatory balance mediated by glia are altered *before* the onset of neurological symptoms. Thus, in forthcoming studies it will be important to study functional alterations in PD microglia in earlier developmental stages using *in vitro* models.

Finally, public datasets have limited diagnostic and clinical information attached to the sequencing data, therefore limiting the interpretations that can be made from sequencing data analysis on clinical outcomes. Due to the nature of the public datasets, we cannot control for factors such as non-demographic patient characteristics, or variance introduced by tissue handling procedures. These effects are inevitable since samples come from a range of biobanks in the USA and the UK and handled by many people before being processed for sequencing.

## 5.0 CONCLUSIONS

SnRNA sequencing data from post-mortem PD patients and matched healthy donors allowed us to profile altered microglial and astrocytic functions in the midbrain regions. An increased expression of chaperone protein binding and protein folding genes suggests a role in clearing protein aggregates by both microglia and astrocytes. Interestingly, a decrease in the expression of inflammation associated genes in microglia implies that later disease stages microglia may be fatigued, thereby performing their normal function with decreased efficacy. Altered expression of genes associated with channel functions in astrocytes suggests that signalling mechanisms may also be disrupted in these supporting cells that often mediate cross-glial communication in the brain. This is supported by the concomitant decreased expression in PD of transmembrane receptor activity genes associated with resting and non-inflammatory activated microglia. However, the limited nature of the datasets prevents us from making a conclusive statement about altered inflammation in PD microglia, since most cytokines and chemokines are expressed in the cytosol or extra-nuclear organelles that are not represented. Future studies will explore functional alterations in microglia and astrocytes in earlier disease stages using *in vitro* models where the dynamic changes in inflammation-mediating cytokines can be tracked over longer periods of time, including biofluid analyses which when paired with sequencing data will reveal a more complete picture of glial functional dynamics.

## 6.0 Availability of data and materials

The original datasets are available publicly through NCBI Gene Expression Omnibus under <u>GSE178265</u> and <u>GSE157783</u>. All scripts used for analysis and data visualization are compiled in a public Github repository, <u>snGlia-PD</u>.

