## Supplementary Figure legend for "Single nuclear RNA sequencing shows altered microglial and astrocytic functions in post-mortem Parkinson’s disease tissue"

**Supplementary Figure legends**

**Supplementary Figure S1. Proportionality of broad cell types in the Kamath dataset.** All cell types are within expected range based on control donors, except for the proportion of dopaminergic neurons (da) which is decreased nearly two-fold in PD derived cells.

**Supplementary Figure S2. Microglial cell extracted from the Smajić dataset show comparable results.** (A) Volcano plot shows altered gene expression in PD microglia with selected genes highlighted. HSP90AB1, HSP90AA1 and GPNMB are also among the top upregulated genes in this set as in Kamath. (B) Top 10 up and downregulated GO molecular function terms in PD microglia (Smajić). (C) CNet plot showing the gene enrichment associated with the top 4 upregulated and top 3 downregulated functions in PD microglia (Smajić).

**Supplementary Figure S3. Astrocytes extracted from Smajić dataset show some consistency with findings in Kamath dataset.** (A) Volcano plot shows altered gene expression in PD astrocytes with selected genes highlighted. (B) GO molecular function results enriched in PD astrocytes (Smajić); due to the minimal gene list resulting from DGE analysis, only a few functions were found to be enriched in this dataset. (C) CNet plot showing the gene enrichment associated with the top 5 altered molecular function terms in PD astrocytes (Smajić).
