## Supplementary figures and images for "Single nuclear RNA sequencing shows altered microglial and astrocytic functions in post-mortem Parkinson’s disease tissue"

### Supplementary Figures

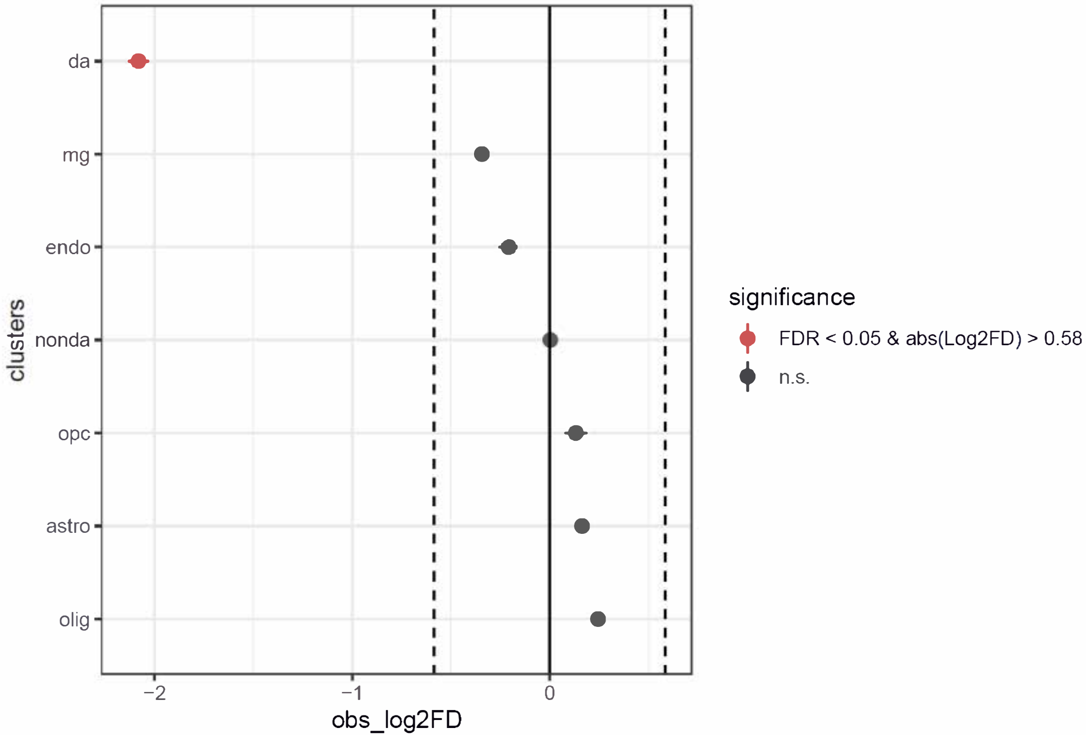


Supplementary Figure S1

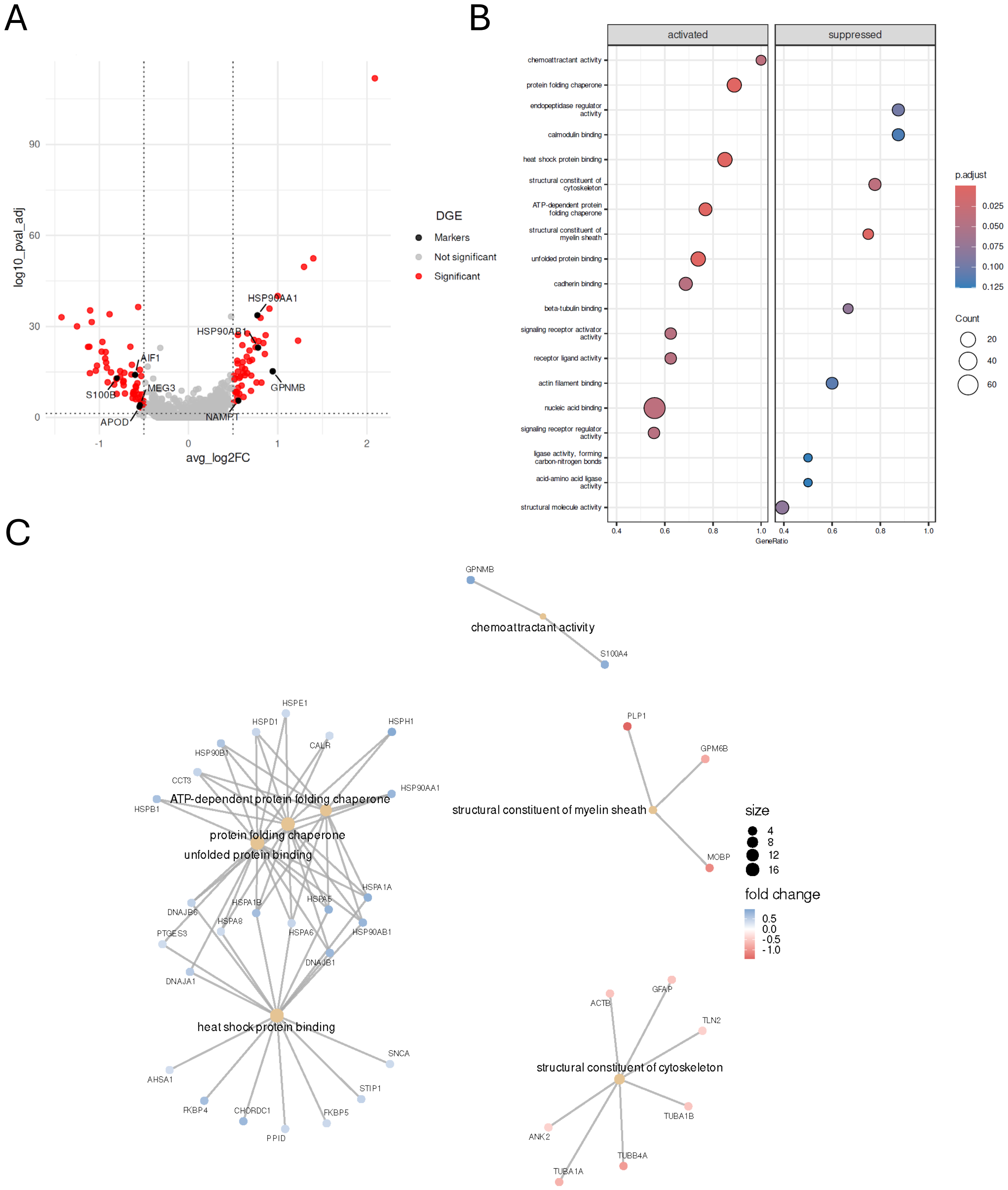


Supplementary Figure S2

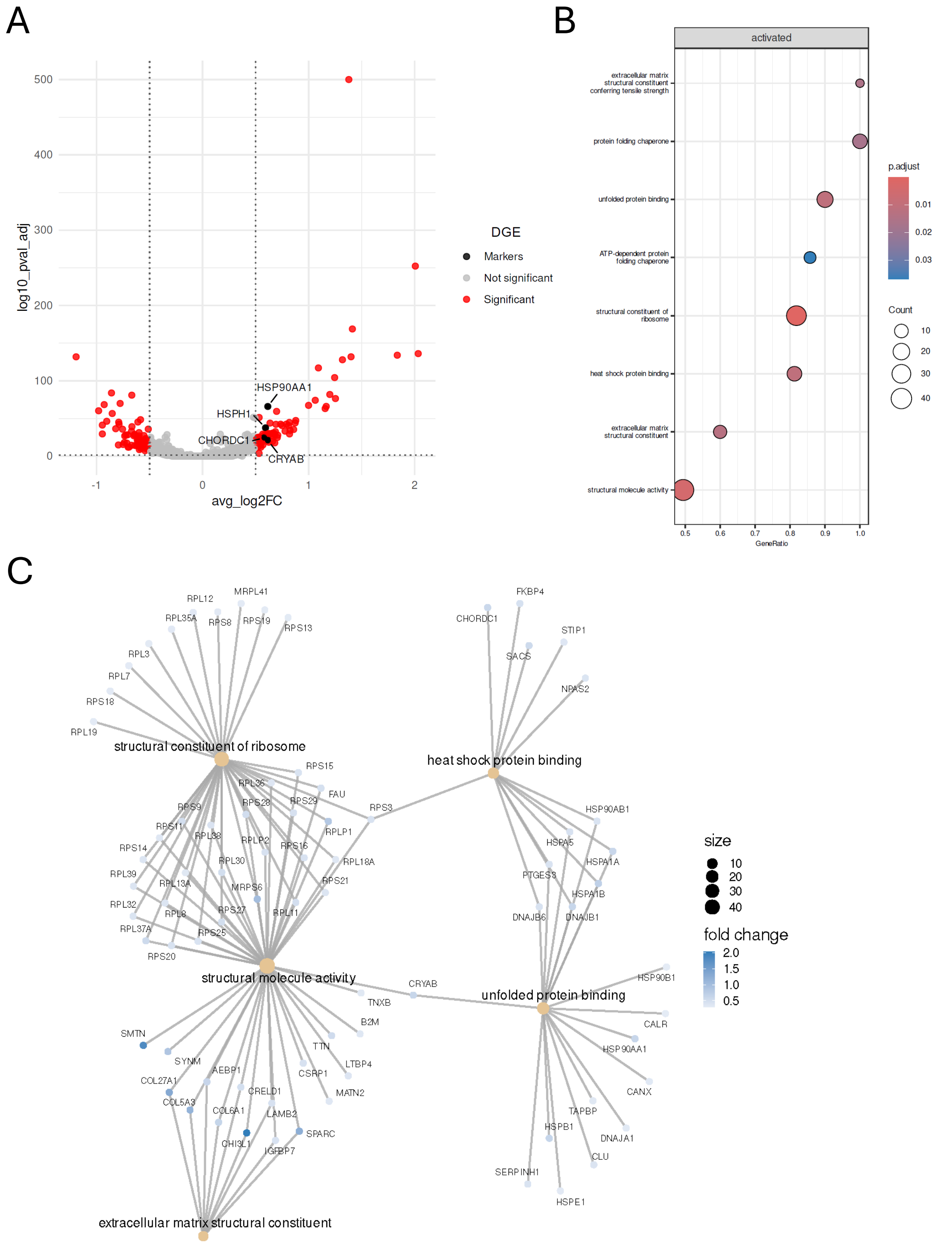


Supplementary Figure S3
